# Oat species and interspecific amphiploids show predominance of diploid nuclei in the syncytial endosperm

**DOI:** 10.1101/2022.12.30.522337

**Authors:** Paulina Tomaszewska, Romuald Kosina

## Abstract

Apart from apomictic types, the *Polygonum*-type eight-nuclear embryo sac is considered to be dominant in grasses. A triploid endosperm is formed as a result of double fertilisation. This study showed, for the first time, the dominance of diploid nuclei in the syncytial stage of the central cell of embryo sac in oat species and amphiploids. The dominance of diploid nuclei, which were the basis for the formation of polyploid nuclei, was weaker in amphiploids due to aneuploid events. The genomic *in situ* hybridisation method applied in the study did not distinguish the maternal and paternal haploid nuclei of embryo sac. However, this method demonstrated the lack of a set of genomes of one haploid nucleus. It can be assumed that the formation of diploid oat endosperm occurred after the fusion of one polar nucleus and the nucleus of a sperm cell, while the second polar nucleus formed 1*n* nuclei. The levels of syncytial ploidy were not influenced by both aneuploid events and correlated with pollen developmental anomalies. The differences in the analysed cytogenetic events distinguished amphiploids and their parental species in the ordination space.

## 1. Introduction

Embryo sacs of Angiospermae constitute a broad spectrum of structural types. The *Polygonum-type* embryo sac occurs in the majority of flowering plants (Willemse and Went, 1984) and is treated almost as a paradigm in the field of botany. It is known that grasses, including cereals, develop this type of embryo sac.

In their study, Friedman and Ryerson (2009) showed that a four-nucleate module can be distinguished in the development of the embryo sac, both in basal angiosperms, such as *Amborella*, and in monocots with the *Polygonum-type*. Finally, the fusion of the polar nuclei from two modules results in the formation of a triploid endosperm. In cereals, a dominant 3C-level polyploidy in a central cell of embryo sac is maintained throughout the development, up to the ripe endosperm tissue, for example in *Zea* or *Hordeum vulgare*. Fluctuations can be seen only between this level and its multiplications 6C, 12C, and 24C (Nguyen et al., 2007; Nowicka et al., 2021). However, the pattern of DNA multiplication was found to remain stable in *Zea mays* hybrids, as it was altered indicating the maternal quantitative effect (Kowles et al., 1997).

In the *Avena* species and amphiploids, Tomaszewska (2017) observed that the embryo sac in the syncytial stage varied from the *Polygonum-type*. Additionally, some deviation from the *Polygonum* development was noted in *Avena fatua*. In addition to the initial 3C and 6C levels, the 2C level appeared in the mature aleurone layer, which is the youngest tissue of the endosperm (Maherchandani and Naylor, 1971). This gives rise to the speculation whether the *‘Polygonum* thinking’ is applicable to cereals, especially the genus *Avena*.

Several detailed studies conducted on *Arabidopsis* have proven that many genes likely have an impact on the development of embryo sac. Certain genes, such as *CKI1* (histidine kinase), can be expressed during this stage, restricting the fusion of the polar nuclei and interacting with the antipodals activity (Li and Yang, 2020). The occurrence of the *Polygonum-type* embryo sac was unstable in an orchid *Maxillaria*, where micropylar and chalazal nuclei remain unfused (Kolomeitseva et al., 2022). This is an example of sexual development and not apomictic development. However, studies show that in *Tulipa* sp. with the tetrasporic *Fritillaria-type* embryo sac and a pentaploid endosperm, the fusion of micropylar (one maternal) and sperm (one paternal) nuclei results in the formation of a diploid endosperm (Mizuochi et al., 2009). These indicate that several variants of embryo sac development can exist.

Apomictic and autonomous development of embryo sac and endosperm, independent of fertilisation, can also be observed in some plants, including grasses (Nogler, 1984; Vinkenoog and Scott, 2001). Apomixis, as a complex developmental process, results in the formation of an endosperm with an altered level of ploidy. A study showed the formation of multiporate pollen grains in apomictic plants (Ma et al., 2009). Such types of pollen grains have been found in the genus *Avena* in artificial amphiploids (Tomaszewska and Kosina, 2022), which suggests the apomictic development of embryo sac.

Natural hybrids and increased ploidy levels are common in plants (Grant, 1981). This phenomenon was also applied to obtain new types of crops, such as Triticale. In both somatic tissues and storage endosperm of plants, it is accompanied by cytogenetic disturbances, which manifest either as amplification or loss of DNA, or even as the elimination of whole genomes, especially in hybrids (Gvaladze et al., 2002; Tomaszewska and Kosina, 2018). In *Avena*, such changes, in particular the high level of translocations with an asymmetric frequency between genomes, results in the differentiation of genomes (Tomaszewska et al., 2022), which highlights the unique cytogenetic status of oats among cereals.

## 2. Material and methods

### 2.1. Plant materials

Species and amphiploids of oats used in the study are listed in Table 1. The investigated *Avena* species were classified based on the nomenclature adopted from https://npgsweb.ars-grin.gov/gringlobal/taxon/taxonomy_search.aspx (accessed on 2 December 2022) and http://www.theplantlist.org/ (accessed on 2 December 2022) and were as follow:

*A. fatua* L.
*A. sterilis* L.
*A. sativa* L.
*A. barbata* Pott ex Link.
*A. abyssinica* Hochst.
*A. magna* H.C. Murphy & Terrell.
*A. strigosa* Schreb.
*A. longiglumis* Durieu.
*A. eriantha* Durieu.

**Table 1.** Accessions of oat amphiploids and parental species used in the study.

| Amphiploids<br>Parentalspecies | Symbols<br>used in<br>figures | Accession<br>number | Donor | Origin | Determined<br>ploidy level<br>in roots | Expected<br>chromosome<br>number in 3n<br>endosperm |
| --- | --- | --- | --- | --- | --- | --- |
| *A. barbata* × *A. sativa* ssp. *nuda* | b/sn | CIav7903 | NSGC | - | octoploid | 84 (12x) |
| *A. eriantha* × *A. sativa* | e/sa | PI458781 | NSGC | - | octoploid | 84 (12x) |
| *A. barbata* × *A. sativa* | b/sa | CIav7901 | NSGC | - | hexaploid | 63 (9x) |
| *A. fatua* × *A. sterilis* | f/ste | CIav9367 | NSGC | - | hexaploid | 63 (9x) |
| *A. magna* × *A. longiglumis* | m/l | CIav9364 | NSGC | - | hexaploid | 63 (9x) |
| *A. abyssinica* × *A. strigosa* | a/str | CIav7423 | NSGC | - | tetraploid | 42 (6x) |
| *A. fatua* L. | Af | - | R. Kosina | Poland | hexaploid | 63 (9x) |
| *A. sterilis* L. | Aste | PI311689 | NSGC | Israel | hexaploid | 63 (9x) |
| *A. sativa* L. | Asa | - | R. Kosina | Poland | hexaploid | 63 (9x) |
| *A. barbata* Pott ex Link | Ab | AVE1938 | BAZ | Spain | tetraploid | 42 (6x) |
| *A. abyssinica* Hochst. | Aa | PI331373 | NSGC | Ethiopia | tetraploid | 42 (6x) |
| *A. magna* Murphy et Terrell | Am | 1786 | VIR | Morocco | tetraploid | 42 (6x) |
| *A. strigosa* Schreb. | Astr | 51624 | BAZ | Belgium | diploid | 21 (3x) |
| *A. longiglumis* Dur. | Al | PI367389 | NSGC | Portugal | diploid | 21 (3x) |
| *A. eriantha* Dur. | Ae | CIav9051 | NSGC | England | diploid | 21 (3x) |

BAZ - Bundesanstalt für Züchtungsforschung an Kulturpflanzen, Braunschweig, Germany; NSGC - National Small Grains Collection, Aberdeen, Idaho, USA; VIR - Vavilov Institute of Plant Industry, St. Petersburg, Russia.

The seeds of oats were germinated for 3 days on a moisturised filter paper at 25°C in the dark. Then, the seedlings were planted into pots filled containing a mixture of soil and sand in a ratio of 3:1. All the plants were grown under the same climatic conditions. Next, young kernels at the syncytial stage of endosperm development were harvested. Based on the morphology of the pistil and the degree of dryness of the stigma, the stage of endosperm development was defined. During five consecutive growing seasons, approximately a total of 200 developing seeds from each accession were collected from the spikelets at 2–4 days after pollination (DAP).

### 2.2. Isolation and fixation of endosperm

Using a binocular microscope, embryo sacs were manually dissected from developing seeds. The peeled embryo sacs were then fixed in 96% ethanol–glacial acetic acid (3:1) fixative for 48 hours at room temperature. This was followed by the fixation of embryo sacs in a freshly prepared fixative solution at 4°C.

### 2.3. Preparation of endosperm chromosomes and genomic in situ hybridisation

Most cytogenetic endosperm analyses have thus far been performed on paraffin sections. However, in this study, the enzymatic maceration method was used to obtain better chromosome spreads. This technique involves squashing chromosomes between a slide and a cover slip by applying thumb pressure. The modified method of Schwarzacher and Heslop-Harrison (2000) was used for this purpose, which was originally developed for chromosome preparation from root tips. The fixed embryo sacs were rinsed in distilled water for 15 minutes, and then washed in 0.01 M citrate buffer three times for 5 min each. Subsequently, the embryo sacs were transferred to a maceration enzyme solution (0.3% cellulase, 0.3% pectolyase, 0.3% cytohelicase in 0.01M citrate buffer) and digested in a hybridisation oven (Biometra OV3) at 37°C for 45 minutes. They were then washed in 0.01 M citrate buffer three times for 5 min each. In the next step, the digested embryo sac was placed on a clean slide in a drop of 45% acetic acid for 1 minute, and then cut open with a scalpel. Only the liquid content of the embryo sac was left on the slide, and the other tissues were discarded. The endosperm tissue was then squashed in 45% acetic acid under a coverslip by applying light thumb pressure. Finally, the slides were frozen in liquid nitrogen and air-dried. Genomic *in situ* hybridisation (GISH) was performed on endosperm as previously described by Tomaszewska and Kosina (2021), using genomic DNAs extracted from fresh leaves of *A. eriantha* (C genome) and *A. nuda* (A genome), labelled with tetramethylrhodamine-5-dUTP (red fluorescence) and digoxigenin-11-dUTP, respectively, as probes. The probe labelled with digoxigenin was detected with fluorescein isothiocyanate-conjugated sheep anti-digoxigenin antibody (green fluorescence). The slides were then counterstained with DAPI and analysed under Olympus BX-50 and BX-60 fluorescence microscopes (Hamburg, Germany). Images were taken with an Olympus E-520 camera (Olympus Imaging Europa GMBH, Hamburg, Germany).

### 2.4 Isolation of pollen grains

Pollen grains were collected during three consecutive vegetation seasons. The frequency of formation of micropollens and multiporate pollens was determined in oat species and amphiploids. Detailed studies on morphotypes and viability of pollen grains in oats have already been published (Tomaszewska and Kosina, 2022).

### 2.5. Correlation and numerical taxonomy analyses

A matrix of correlation coefficients and data for nonmetric multidimensional scaling (nmMDS) were obtained using Rohlf’s approach (Rohlf, 1994).

## 3. Results and Discussion

### 3.1. Variation of chromosome number

The chromosome number variation in the cereal endosperm was larger than that observed in other vegetative tissues, which can be due to significant DNA amplification and aneuploidy events (Kaltsikes and Roupakias, 1975; Nowicka et al., 2021). In the oat endosperm, amphiploids originated from hexaploid parents showed a larger variation compared to those having 2*x* and 4*x* parents. Among the oat species, *A.magna* (4*x*) and *A*. *eriantha* (2*x*) showed extreme variation in chromosome numbers. It was observed that amphiploids had a twofold larger range of chromosome numbers than parental species. The position of modal values for the 2*n* chromosome numbers indicates the left-hand skewness of distributions in most taxa. The only exceptions were *A*. *magna* (reverse-J shaped distribution) and *A*. *longiglumis* and *A*. *eriantha* (right-skewed distribution), while some showed nearly symmetric distributions (Table 2).

**Table 2.** Number of chromosomes observed in the syncytial endosperm of oat species and amphiploids.

| Amphiploids / parental species | Number of analysed metaphases | Number of chromosomes | rcn |
| --- | --- | --- | --- |
| *A. barbata* × *A. sativa* ssp. *nuda* | 179 | 35, 36, 41, 42, 47, 51, 52, 54, 55, **56**, 57, 61, 104, hp | 69 |
| *A. eriantha* × *A. sativa* | 201 | 28, 40, 42, 48, 49, 50, 52, 53, 54, 55, **56**, 104 | 76 |
| *A. barbata* × *A. sativa* | 85 | 21, 32, 36, 37, 38, **42**, 44, 48, 56, 63, hp | 42 |
| *A. fatua* × *A. sterilis* | 203 | 25, 28, 30, 32, 35, 37, 38, 39, 40, 41, **42**, 61, 63, 84, hp | 59 |
| *A. magna* × *A. longiglumis* | 116 | 39, 40, 41, **42**, 52, 63, hp | 24 |
| *A. abyssinica* × *A. strigosa* | 164 | 14, 19, 21, 26, 27, **28**, 42, hp | 28 |
| Σ/n (for amphiploids) |  |  | 49.7 |
| *A. fatua* | 174 | 21, 26, 28, 37, 40, 41+1r, **42**, 56 | 39 |
| *A. sterilis* | 141 | 41, **42**, 78, 84 | 43 |
| *A. sativa* | 91 | 36, 38, 40, **42** | 6 |
| *A. barbata* | 91 | 26, **28**, 42 | 16 |
| *A. abyssinica* | 68 | 13, 14, 26, 27, **28** | 15 |
| *A. magna* | 111 | **28**, 42 | 14 |
| *A. strigosa* | 194 | 10, 11, 12, 13, **14**, 17, 19, 21, 23, 28, hp | 18 |
| *A. longiglumis* | 49 | 12, **14**, 21, 28, 36, 38, 42, hp | 30 |
| *A. eriantha* | 203 | 7, 12, 13, **14**, 14+1r, 17+1r, 18+1r, | 35 |
| Σ/n (for species) |  |  | 24.0 |

The development of a *Polygonum-type* embryo sac with a triploid endosperm is common in Angiospermae. DNA amplification runs from 3C through 6C and 12C up to 24C in grasses with their cultivated taxa, such as wheat or barley (Chojecki et al., 1986; Nowicka et al., 2021). There were much higher levels of ploidy in rice and sorghum, rising to over 96C in maize (Nguyen et al., 2007). In Table 2, a high level of nuclei ploidy is indicated by the symbol ‘hp’, and the detailed values hidden therein would undoubtedly widen the range of chromosome numbers. In Table 3, the series of chromosome numbers refer to aneuploid states, which were more numerous in amphiploids than in species. Unlike other cereals, endosperm cytogenetic variability is greater in the genus *Avena*, both in hybrids and in species. This may be due to the increase in cytogenetic variability by aneuploidy. The fusion of two smaller nuclei, among others, can result in the formation of hp-type nuclei with a high and variable number of chromosomes. This is evidenced by the formation of giant nuclei in *Arabidopsis* (Baroux et al., 2004), and dumbbell nuclei in Triticale (Kaltsikes and Roupakias, 1975), as well as in *Avena* species and amphiploids (Tomaszewska, 2017). Furthermore, the reproductive system of the plant changes from generative to apomictic at very high levels of ploidy. The *Bouteloua curtipendula* grass complex with basic chromosome number *x* = 10 is an example of the above, in which agamic taxa are formed at the borderline level of 2*n* = 52 (Grant, 1981).

**Table 3.** Percentage of metaphases in the syncytial endosperm of different ploidy level.

| Amphiploid / parental species | 1n | 2n | 3n | 4n | 6n | ap | hp |
| --- | --- | --- | --- | --- | --- | --- | --- |
| *A. barbata* × *A. sativa* ssp. *nuda* | 0 | 57.14 | 0 | 0 | 0 | 40.82 | 2.04 |
| *A. eriantha* × *A. sativa* | 1.96 | 54.90 | 0 | 0 | 0 | 43.14 | 0 |
| *A. barbata* × *A. sativa* | 2.17 | 69.57 | 2.17 | 0 | 0 | 23.92 | 2.17 |
| *A. fatua* × *A. sterilis* | 1.04 | 64.58 | 2.08 | 1.04 | 0 | 30.22 | 1.04 |
| *A. magna* × *A. longiglumis* | 0 | 88.52 | 1.64 | 0 | 0 | 8.20 | 1.64 |
| *A. abyssinica* × *A. strigosa* | 1.64 | 59.02 | 14.75 | 0 | 0 | 22.95 | 1.64 |
| Σ/n (for amphiploids) | **1.14** | **65.62** | **3.44** | **0.17** | **0** | **28.21** | **1.42** |
| *A. fatua* | 6.52 | 80.43 | 0 | 0 | 0 | 13.05 | 0 |
| *A. sterilis* | 0 | 87.18 | 0 | 2.56 | 0 | 10.26 | 0 |
| *A. sativa* | 0 | 90.00 | 0 | 0 | 0 | 10.00 | 0 |
| *A. barbata* | 0 | 70.83 | 20.83 | 0 | 0 | 8.34 | 0 |
| *A. abyssinica* | 14.29 | 71.43 | 0 | 0 | 0 | 14.28 | 0 |
| *A. magna* | 0 | 95.00 | 5.00 | 0 | 0 | 0 | 0 |
| *A. strigosa* | 0 | 66.06 | 24.24 | 1.21 | 0 | 7.88 | 0.61 |
| *A. longiglumis* | 0 | 29.03 | 25.81 | 9.68 | 22.58 | 9.67 | 3.23 |
| *A. eriantha* | 0.68 | 71.92 | 17.12 | 0.68 | 0.68 | 8.24 | 0.68 |
| Σ/n (for species) | **2.39** | **73.54** | **10.33** | **1.57** | **2.58** | **9.08** | **0.50** |

1n...6n – levels of ploidy, ap – aneuploidy, hp – hyperploidy

### 3.2. Ploidy levels of syncytium

All the studied taxa showed a dominance of diploid nuclei (2*n*) in the syncytial endosperm, which was more frequent in the parental species (Table 3; Fig. 1a–f). Therefore, the phenomenon of diploid syncytium (and cellular endosperm?) can be assumed for the whole *Avena* genus. Two species, *A. magna* and *A*. *longiglumis*, showed the greatest difference of dominance, with 95% of diploid nuclei and only 29.03%, respectively. Triploid nuclei were not numerous in amphiploids but were more frequent in tetraploid (*A*. *barbata*) and diploids (*A*. *strigosa, A. longiglumis*, and *A*. *eriantha*). Some uniformity of nuclei frequency was observed at higher levels of ploidy, as in the case of *A*. *longiglumis*, which were formed at the cost of the 2*n* nuclei. Hyperploidy and aneuploidy were threefold more frequent in amphiploids than in parental species. The highly polyploid nuclei were absent in 4*n* and 6*n* species. It was noted that *A*. *abyssinica* and *A*. *fatua* had a high percentage of 1*n* nuclei. No 3*n* nuclei have been recorded in these species, because they would just be formed from the union of 1*n* and 2*n*. The differences between taxa in the absence or presence of 4*n* or 6*n* nuclei indicate the intertaxa variation of the polyploidisation rate. The GISH method enabled the detection of A and C genomes, which showed a lack of genomes of one haploid nucleus (Fig. 1a, b). This suggests that one nucleus, maternal polar or sperm male, did not fuse to form the triploid nucleus of the central cell. It should be noted that the GISH method does not allow to distinguish female polar nuclei from male sperm nuclei as they are genomically identical.

**Fig. 1.**
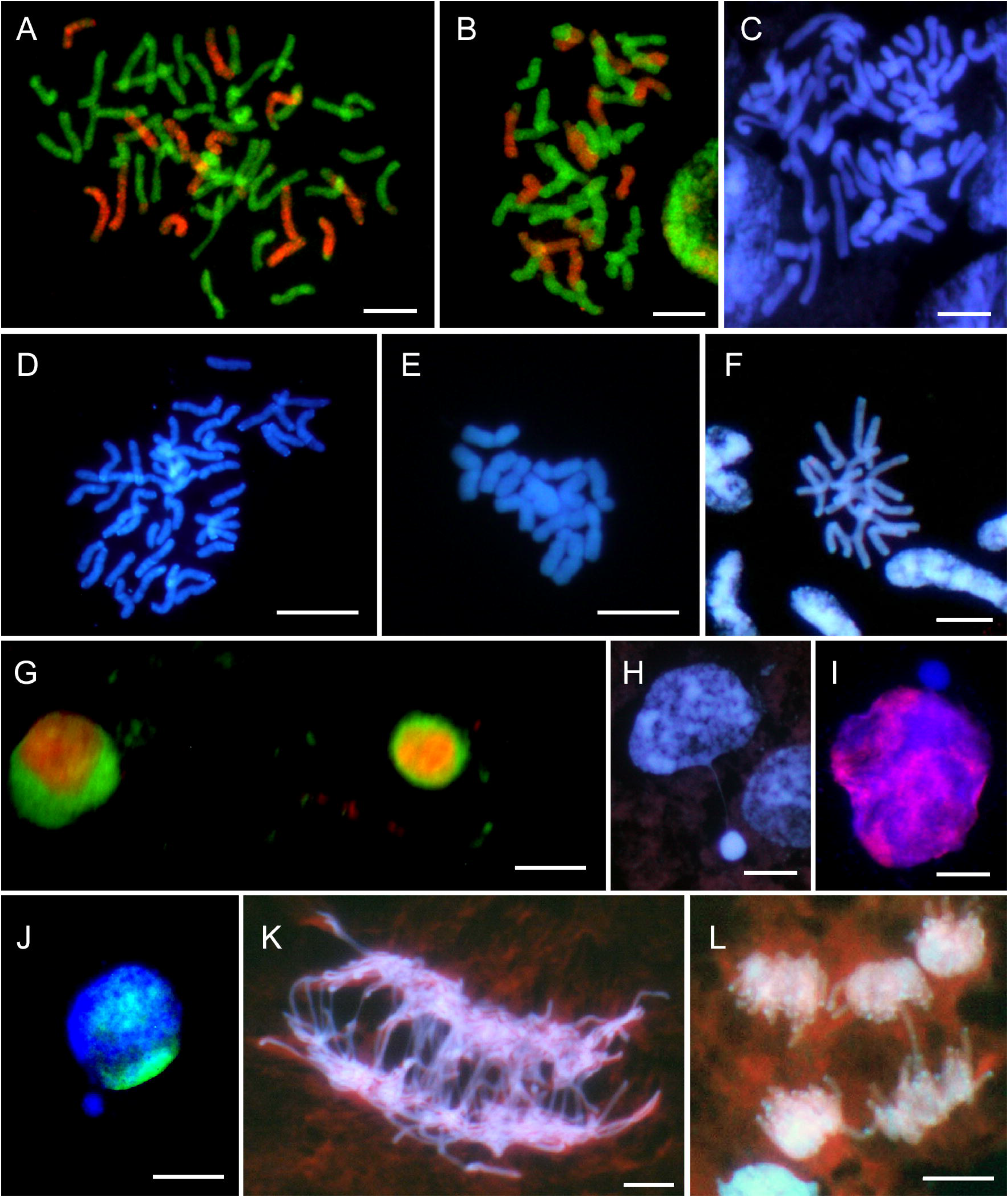
Data on 2*n* chromosome numbers and some cytogenetic disorders in the oat syncytial endosperm. **A** - GISH metaphase in *A. barbata* × *A. sativa* ssp. nuda with 2*n* = 56; 14 red chromosomes of genome C, 42 green chromosomes of genomes A and D; there is a lack of one genome C and 21 chromosomes of A/B/D genomes. **B** - GISH metaphase in *A. barbata* × *A. sativa* with 2*n* = 42; 14 red chromosomes of genome C, 28 green chromosomes of genomes A and D; there is a lack of one genome C and 14 chromosomes of A/B/D genomes. **C** - DAPI metaphase in *A. eriantha* × *A. sativa* with 2*n* = 56. **D** -DAPI metaphase in *A. fatua* × *A. sterilis* with 2*n* = 42. **E** - DAPI metaphase in *A. strigosa* with 2*n* = 14. **F** - DAPI metaphase in *A. eriantha* with 2*n* = 14. **G** - GISH interphase nuclei in *A. sterilis* with green external A/D genomes and internal red C genome. **H** - DAPI prophase nuclei with condensed micronucleus connected by a thread of chromatin with upper left nucleus in *A. eriantha* × *A. sativa*. **I** - An early GISH prophase nucleus with an extruded micronucleus composed of A/D genomes in blue (genomes C, red/violet) in *A. barbata* × *A. sativa* ssp. nuda. **J** - A GISH interphase-prophase nucleus in *A. abyssinica* with an extruded micronucleus composed of B genome (A genome in green). **K** - DAPI anaphase with multiple bridges in *A. barbata* × *A. sativa*. **L** - DAPI multipolar anaphases in *A. eriantha*. Scale bars = 10 μm

When the polar nuclei of the central cell do not merge, one of them fuses with the nucleus of the sperm cell, while the other remains free and divides giving rise to nuclei 1*n*. The nucleus of the second sperm cell fuses with that of the egg cell forming a zygote. This results in the diploid level of the syncytial endosperm of oats. The fusion of two polar nuclei into a secondary nucleus, without the participation of a sperm nucleus, may be another possible version of nuclear fusion. In this case, the sperm nucleus remains free, divides, and produces 1*n*-type nuclei. However, this is less probable due to the nonequivalent functioning of the free polar nucleus compared to the free sperm nucleus. The free sperm nucleus enters the central cell with degraded mitochondria and their mtDNA (Sato and Sato, 2013). It is well-known that mitochondria play an important role in nuclear fusion (Li and Yang, 2020). Finally, the sperm nucleus will be degraded, and the central cell will develop as the maternal autonomous-type endosperm.

In this study, the level of pollen viability was higher than 90% and pollen development anomalies were at a level lower than 1% in all the investigated *Avena* accessions. The development of anomalous, multiporate pollen grains was observed only in the case of amphiploids (Tomaszewska and Kosina, 2022). Plant development and seed setting in the field cultivation conditions was found to be normal (R. Kosina, unpubl.). Such vigor of offspring indicates generative reproduction with the involvement of both nuclei of sperm cells. Thus, the normal double fertilisation of the secondary diploid nucleus is related only to one haploid nucleus of the central cell of the embryo sac, and therefore, differs from the *Polygonum*-type. In *Arabidopsis* mutants, the lack of the fusion of polar nuclei in the central cell was shown to be due to the activity of genes such as *GFA2, AAC2, SYCO/FIONA, GCD1, CLO, MAA3, Bip1/2, SEC22*, and *CKI1* (Li and Yang, 2020). The *fie* (*fertilization-independent endosperm*) mutation analysed in the development of pods and seeds in *Arabidopsis* (Ohad et al., 1996) and the *fis2* mutation (Chaudhury et al., 2001) indicated the fusion of the polar nuclei without the participation of the sperm nucleus. Such endosperm is of the maternal, autonomous type. The development of endosperm did not surpass the syncytial stage in the absence of pollination in the *fie* mutation. This independence of the development of endosperm, in which numerous genes are imprinted, can be attributed to the changes in the imprinting and activation of genes (Vinkenoog and Scott, 2001). As observed in the studies of apomictic grasses (Ma et al., 2009), the development of multiporate pollen grains in the studied amphiploids (Tomaszewska and Kosina, 2022) may indicate apomictic reproduction in oats. This feature was noted only in amphiploids among the examined oats, while the domination of the diploid syncytium was observed in the entire group, particularly in the species. Therefore, it can be assumed that anomalous multiporate pollen grains lose the competition with normal pollens due to the slow growth of numerous pollen tubes. Further, material collected from a pool of embryo sacs was analysed on each microscope slide. Apart from common in plants pseudogamy (Nogler, 1984; Chaudhury et al., 2001), another type of apomictic development cannot be ruled out for a single embryo sac in oat amphiploids with a higher frequency of multiporate pollen grains.

The cytogenetic status of the syncytial endosperm was highly variable as well, in addition to the dominance of diploid nuclei. It was observed that different events increased the frequency of aneuploidy. Concentric (Fig. 1g) or side-by-side spatial separation of genomes likely facilitated the elimination of monogenomic micronuclei (Fig. 1i, j). Furthermore, micronuclei with mixed genome composition were also eliminated (Tomaszewska and Kosina, 2021). The micronuclei generally showed a high degree of chromatin condensation (Fig. 1h). The occurrence of multiple bridges (Fig. 1k) and elimination of single chromosomes or their fragments were very common. Additionally, there were only a few multipolar anaphases (Fig. 1l).

### 3.3. Correlation analysis

Table 4 shows the results of the correlation analysis of the characters with the addition of two other traits describing anomalous pollen types (see Table 3). The analysis revealed two sets of significantly correlated traits. These traits were related to the ploidy levels of nuclei and the frequencies of anomalous nuclei and the disturbances in the development of pollen grains. The correlation analysis proved that the diploid level of syncytial nuclei was not influenced by anomalous mitoses that produce aneuploid nuclei in endosperm and the increased range of chromosome numbers. The negative coefficients of correlation between the frequency of 2*n* nuclei and the higher ploidy ones (4*n*, 6*n*, hp) indicated that the first group of nuclei changed into the second. The highest coefficient of correlation 0.96*** between the frequency of 4*n* nuclei and that of 6*n* nuclei showed the highest rate of formation of these nuclei from the 2*n* nuclei. The presence of 6*n* nuclei was only observed in a few of the studied taxa (Table 3); however, it can be increased at later stages of endosperm development. The high frequency of aneuploidy and the range of the chromosome numbers correlated with anomalous microsporogenesis, and such interrelations were more frequent in amphiploids (see Tables 2 and 3).

**Table 4.** Pearson’s correlation coefficients matrix of the syncytial endosperm cytogenetic and pollen grain traits (*n* = 15, pooled sample of species and amphiploids).

| Traits | 1n | 2n | 3n | 4n | 6n | hp | ap | mp | mpp | rcn |
| --- | --- | --- | --- | --- | --- | --- | --- | --- | --- | --- |
| 1n | 1.00 |  |  |  |  |  |  |  |  |  |
| 2n | 0.03 | 1.00 |  |  |  |  |  |  |  |  |
| 3n | -0.32 | -0.50* | 1.00 |  |  |  |  |  |  |  |
| 4n | -0.20 | -0.62** | 0.51* | 1.00 |  |  |  |  |  |  |
| 6n | -0.14 | -0.68** | 0.52* | 0.96*** | 1.00 |  |  |  |  |  |
| hp | -0.28 | -0.64** | 0.31 | 0.56* | 0.63** | 1.00 |  |  |  |  |
| ap | 0.04 | -0.47 | -0.38 | -0.20 | -0.16 | 0.22 | 1.00 |  |  |  |
| mp | 0.12 | -0.25 | -0.47 | -0.21 | -0.25 | 0.05 | 0.81*** | 1.00 |  |  |
| mpp | -0.16 | -0.28 | -0.29 | -0.16 | -0.12 | 0.31 | 0.70** | 0.73*** | 1.00 |  |
| rcn | -0.12 | -0.41 | -0.35 | -0.02 | -0.06 | 0.21 | 0.86*** | 0.76*** | 0.63** | 1.00 |

1n...6n – levels of ploidy; hp – hyperploidy; ap – aneuploidy; mp – micropollens; mpp – multiporate pollens; rcn – range of chromosome number; \*, \*\*, \*\*\*—significance levels at $\alpha = 0.05, 0.01, 0.001$ , respectively

### 3.4. Ordination of taxa in the minimum spanning tree

The species *A. longiglumis* was situated at the extreme point of the ordination space (maximum values of *x* and *y* axes) (Fig. 2). This position was determined by the frequencies of nuclei at all the examined ploidy levels. As a rule, parents showing larger values of ordination axes were well distinguished from amphiploids with smaller values. However, the amphiploid *A. magna* × *A. longiglumis* deviated from this rule, which showed the highest frequency of 2*n* nuclei and the lowest frequency of aneuploid nuclei. This agrees with the data presented in Tables 2 and 3, when a matrix of Euclidean distances between taxa was used as a primary set to put operational taxonomic units (OTUs) in the ordination space.

**Fig. 2.**
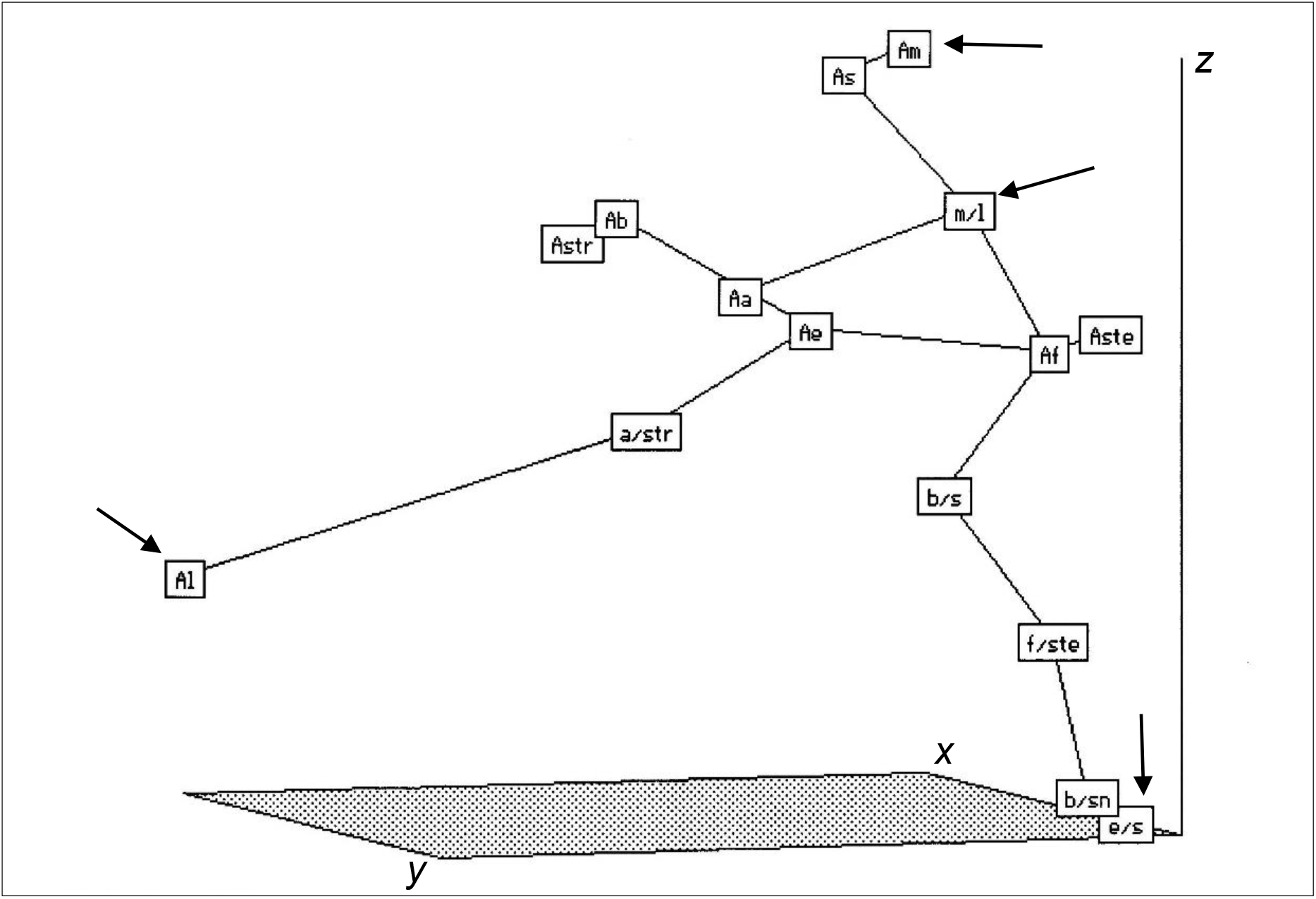
Minimum spanning tree (MST) of amphiploids and parental species (Operational Taxonomic Units, OTUs) of the genus *Avena* in the ordination space (*x*, *y*,and *z* axes) created using Kruskal’s nmMDS method (Rohlf, 1994). The OTUs were described by 10 characters presented in Table 4. Abbreviations of OTUs are given in Table 1. The extreme distances between the pairs of OTUs as shown by the MST are: maximum - a/str-Al **1.194** and minimum - Ab-Astr **0.144**. Some extreme OTUs are marked with arrows.

The numerical taxonomy methods used showed clear discrimination between the oat species and the amphiploids in the structural characters of the mature endosperm (Tomaszewska and Kosina, 2018) and the features of pollen grains (Tomaszewska and Kosina, 2022). These methods also indicated weaker discrimination for the characters listed in Table 4, as these characters describe two different types of events, which were separated in the correlation coefficient matrix. The extreme locations of two species, *A. magna* and *A. longiglumis*, in the ordination space are justified by their characteristics (distributions) in Tables 2 and 3. Data in Table 4 indicate the possible elimination of defective nuclei in the syncytial endosperm, regardless of the ploidy level. Kosina (2016), in their study, pointed to the universality of such elimination. This is due to the high variability of the nuclei in the syncytium. The study by Kaltsikes and Roupakias (1975) indicated extreme variability in nuclei shapes in wheat-rye addition and substitution lines. Another study by Bannikova (1975) revealed excellent embryological characteristics with several anomalies in nuclei shapes and the number of nucleoi in a central cell of a hybrid *Hordeum vulgare* × *Secale cereale*. Additionally, the author documented the presence of irregular nuclei, and their condensation and elimination in syncytial endosperm and initially cellularised endosperm in a hybrid *Triticum durum* × *Secale cereale*. The variability of chromosome numbers and ploidy narrowed during the maturation of the endosperm and the elimination of defective structures. The distributions describing the above variability can show platycurtosis (flattening) (Nogler, 1984). These show that the development of the embryo and endosperm in the genus *Avena* differs from that of the *Polygonum*-type. Compared to the triploid endosperm, the diploid endosperm has a greater energy potential use in karyokinesis (lower number of chromosomes) and cytokinesis. It is uncertain whether the *Avena* diploid syncytium can transform into a diploid cellular endosperm. In *H. vulgare*, the 3C and 6C nuclei dominated during the syncytium stage of endosperm development, and this condition persisted until the maturity of caryopsis (Nowicka et al., 2021). Other nuclei (12C, 24C), which are the multiples of 3C and 6C, had a lower frequency. These findings demonstrate the stability of the main ploidal level of the endosperm. However, the analyses of DNA content in young cells of the aleurone layer in *A. fatua* confirmed the levels of 3C and 6C, as well as the presence of 2C nuclei among nuclei within a large range of DNA content at the stage of full maturity of the aleurone layer (Maherchandani and Naylor, 1971). Among the species listed in Table 3, *A. fatua* has a higher frequency of 1*n* and aneuploid nuclei. This explains the variation in DNA levels to a certain extent, but the appearance of 2C nuclei in the aleurone layer is interesting.

## 4. Conclusions

It has been shown that the chromosome numbers in the syncytial endosperm greatly varies for a wide range of oats, species, and amphiploids. This is also reflected in the shapes of the trait frequency distributions. However, diploid nuclei are the most dominant group. This cytogenetic pattern is common regardless of the level of ploidy of species and diversity of amphiploids. Therefore, it can be assumed that this applies to the whole *Avena* genus. Diploid syncytium, as well as cellular endosperm later, may possibly arise when the polar nuclei of the central cell are not fused. The 1*n* nuclei likely arise from a free polar nucleus, since a defective sperm cell (mtDNA) and its nucleus cannot play such a role. The fusion of dumbbell nuclei with a lower DNA content results in the formation of nuclei with higher levels of ploidy. Defective structures, such as nuclei, chromosomes, their fragments, and micronuclei, with a different DNA content than 1*n* and its multiples, increase the frequency of aneuploidy. The phenomenon of aneuploidy does not correlate with a basic syncytial ploidy, but correlates with anomalous pollen development. The dominance of diploid nuclei in the oat endosperm can be thoroughly understood by performing advanced embryological analyses. The results presented in this paper show that the oat endosperm does not develop according to the *Polygonum-type* embryo sac but its modification.

## Author contribution

PT designed and conducted cytogenetic analyses, interpreted the results and discovered 2*n* syncytium phenomenon. RK provided correlation and numerical data, and supervised the experiments and research. PT and RK prepared figures and wrote the article. The authors read and approved the manuscript.

## Acknowledgements

The authors would like to thank the following institutions for their generous provision of seeds: National Small Grains Collection (Aberdeen, Idaho, USA), Vavilov Institute of Plant Industry (St. Petersburg, Russia), and Bundesanstalt für Züchtungsforschung an Kulturpflanzen (Braunschweig, Germany).

## References

Bannikova, V.P., 1975. Citoèmbriologiâ mežvidovojne sovmestimosti u rastenij. NaukovaDumka, Kiev.

Baroux, C., Fransz, P., Grossniklaus, U., 2004. Nuclear fusions contribute to polyploidization of the gigantic nuclei in the chalazal endosperm of *Arabidopsis*. Planta 220, 38–46. doi: 10.1007/s00425-004-1326-2

Chaudhury, A.M., Ming, L., Miller, C., Craig, S., Dennis, E.S., Peacock, W.J., 1997. Fertilization independent seed development in *Arabidopsis thaliana*. Proc. Natl. Acad. Sci. USA 94, 4223–28.

Chojecki A.J.S., Bayliss, M.W., Gale, M.D., 1986. Cell production and DNA accumulation in the wheat endosperm, and their association with grain weight. Ann. Bot. 58, 809–817.

Friedman W.E., Ryerson, F.K., 2009. Reconstructing the ancestral female gametophyte of angiosperms: insights from *Amborella* and other ancient lineages of flowering plants. Am. J. Bot. 96, 129–143.

Grant V., 1981. Plant speciation. Columbia University Press, New York.

Gvaladze, G., Nadirashvili, N., Akhalkatsi M., 2002. Chromatin diminution during endosperm development in *Allium atroviolaceum* Boiss. (Alliaceae). Bull. Georg. Acad. Sci. 166, 537–540.

Kaltsikes, P.J., Roupakias, D.G., 1975. Endosperm abnormalities in *Triticum-Secale* combinations. II. Addition and substitution lines. Can. J. Bot. 53, 2068–2076.

Kolomeitseva, G.L., Babosha, A.V., Ryabchenko, A.S., 2022. Megasporogenesis, megagametogenesis, and embryogenesis in *Maxillaria crassifolia* (Lindl.) Rchb.f. (Cymbidieae, Orchidaceae). Protoplasma259, 885–903. doi: 10.1007/s00709-021-01710-5

Kosina, R., 2016. Grass endospermal nuclei are selected by means of apoptosis. Annu. Wheat Newsl. 62, 46–47.

Kowles, R.V., Yerk, G.L., Haas, K.M., Phillips R.L., 1997. Maternal effects influencing DNA endoreduplication in developing endosperm of *Zea mays*. Genome 40, 798–805. doi: 10.1139/g97-803

Li, H.-J., Yang, W.-C., 2020. Central cell in flowering plants: specification, signaling, and evolution. Front. Plant Sci. 11: 590307. doi: 10.3389/fpls.2020.590307.

Ma, G., Huang, X., Xu, Q., Bunn, E., 2009. Multiporate pollen and apomixis in Panicoideae. Pak. J. Bot. 41, 2073–2082.

Maherchandani, N.J., Naylor, J.M., 1971. Variability in DNA content and nuclear morphology of the aleurone cells of *Avena fatua* (wild oats). Can. J. Genet. Cytol. 13, 578–584.

Mizuochi, H., Matsuzaki, H., Moue, T. et al., 2009. Diploid endosperm formation in *Tulipa* spp. and identification of a 1:1 maternal-to-paternal genome ratio in endosperms of *T. gesneriana* L. Sex. Plant Reprod. 22, 27–36. doi: 10.1007/s00497-008-0088-6

Nguyen H.N., Sabelli P.A., Larkins B.A., 2007. Endoreduplication and programmed cell death in the cereal endosperm. In: Olsen, O.-A. (Ed.), Endosperm - developmental and molecular biology. Plant Cell Monograph 8. Springer Verlag, Berlin, Heidelberg, pp. 21–43. doi: 10.1007/7089_2007_106

Nogler, G.A., 1984. Gametophyticapomixis. In: Johri B.M., (Ed.) Embryology of Angiosperms. Springer, Berlin, Heidelberg, pp. 475–518.

Nowicka, A., Kovacik, M., Tokarz, B., Vrána, J., Zhang, Y., Weigt, D., Doležel, J., Pecinka, A., 2021. Dynamics of endoreduplication in developing barley seeds. J. Exp. Bot. 72, 268–282. doi: 10.1093/jxb/eraa453

Ohad, N., Margossian, L., Hsu, D.-C., Williams, C., Repetti, P., Fischer, R.L., 1996. A mutation that allows endosperm development without fertilization. Proc. Natl. Acad. Sci. USA 93, 5319–5324.

Sato, M., Sato, K., 2013. Maternal inheritance of mitochondrial DNA by diverse mechanisms to eliminate paternal mitochondrial DNA. Biochim. Biophys. Acta (BBA)-Molecular Cell 1833, 1979–1984. doi: 10.1016/j.bbamcr.2013.03.010

Rohlf, F.J., 1994. NTSYS-pc. Numerical taxonomy and multivariate analysis system. Version 1.80. Exeter Software, New York.

Schwarzacher, T., Heslop-Harrison, J.S., 2000. Practical in situ hybridization. Oxford Bios.

Tomaszewska, P., 2017. Mikrostrukturalna i cytogenetyczna analiza bielma wybranych międzygatunkowych amfiploidów rodzaju *Avena* L. Dissertation, University of Wrocław.

Tomaszewska, P., Kosina, R., 2018. Instability of endosperm development in amphiploids and their parental species in the genus *Avena* L. Plant Cell. Rep. 37, 1145–1158. doi: 10.1007/s00299-018-2301-x

Tomaszewska, P., Kosina, R., 2021. Cytogenetic events in the endosperm of amphiploid *Avena magna* × *A. longiglumis*. J. Plant Res. 134, 1047–1060. doi: 10.1007/s10265-021-01314-3

Tomaszewska, P., Kosina, R., 2022. Variability in the quality of pollen grains in oat amphiploids and their parental species. Braz. J. Bot. 45, 987–1000. doi: 10.1007/s40415-022-00822-3

Tomaszewska, P., Schwarzacher, T., Heslop-Harrison, J.S.(P), 2022. Oat chromosome and genome evolution defined by widespread terminal intergenomic translocations in polyploids. Front. Plant Sci. 13:1026364. doi: 10.3389/fpls.2022.1026364

Vinkenoog, R., Scott R.J., 2001. Autonomous endosperm development in flowering plants: how to overcome the imprinting problem? Sex. Plant. Reprod. 14, 189–194.

Willemse, M.T.M., Went, J.L. van, 1984. The female gametophyte. In: Johri B.M., (Ed.) Embryology of Angiosperms. Springer, Berlin, Heidelberg, pp. 159–196.

